# The histopathological staging of tau, but not amyloid, corresponds to antemortem cognitive status, dementia stage, functional abilities, and neuropsychiatric symptoms

**DOI:** 10.1101/671651

**Authors:** Charles B Malpas, Sifat Sharmin, Tomas Kalincik

**Author notes:** Corresponding author: Charles B Malpas, PhD, Clinical Outcomes Research Unit (CORe), Department of Medicine, Royal Melbourne Hospital, The University of Melbourne, 3010, Melbourne, VIC, Australia.

## Abstract

**Objective:** Alzheimer’s disease (AD) is characterised by two cardinal pathologies, namely the extracellular accumulation amyloid-related aggregates, and the intracellular formation of tau-related neurofibrillary tangles (NFTs). While both pathologies disrupt cognitive function, a large body of evidence suggests that tau-pathology has a stronger relationship with the clinical manifestation of the disease compared to amyloid. Given the ordinal nature of histopathological staging systems, however, it is possible that the effect of amyloid pathology has been underestimated in clinicopathological studies.

**Method:** We investigated this possibility using data from the National Alzheimer’s Coordinating Center (NACC) database. Bayesian ordinal models were used to directly investigate the relative contribution of Braak NFT, diffuse plaque, and neuritic plaque staging to the severity of antemortem clinical impairment.

**Results:** Data from 144 participants were included in the final analysis. Bayesian ordinal models revealed that Braak NFT stage was the only predictor of global cognitive status, clinical dementia stage, functional abilities, and neuropsychiatric symptoms. When compared directly, Braak NFT stage was a stronger predictor than diffuse or neuritic plaques across these domains.

**Conclusions:** These findings confirm that tau-related pathology is more strongly related to clinical status than amyloid pathology. This suggests that conventional clinical markers of disease progression might be insensitive to amyloid-pathology, and hence might be inappropriate for use as outcome measures in therapeutic trials that directly target amyloid.

## Introduction

Alzheimer’s disease (AD) is characterised by two cardinal pathologies, namely the extracellular accumulation amyloid-related aggregates, and the intracellular formation of tau-related neurofibrillary tangles (Haass & Selkoe, 2007). Both of these pathologies disrupt normal neuronal function but develop according to different anatomical trajectories (Arnold, Hyman, Flory, Damasio, & Van Hoesen, 1991; Arriagada, Growdon, Hedley-Whyte, & Hyman, 1992). While cerebral tau-pathology first forms in mesial temporal structures, amyloid-pathology first accumulates in neocortex (Braak, Alafuzoff, Arzberger, Kretzschmar, & Del Tredici, 2006; Braak & Braak, 1991; Thal, Rüb, Orantes, & Braak, 2002). The eventual distribution of these pathologies reflects their origins. The mesial temporal structures are the most heavily affected by tau-pathology while amyloid-pathology is most prominently deposited in neocortical regions (Cho et al., 2016). This raises the possibility that the cardinal pathologies preferentially disrupt different brain networks, resulting in differential contributions to the cognitive expression of AD (Malpas, Saling, Velakoulis, Desmond, & O’Brien, 2015).

A large body of research has addressed the pathological basis of cognitive impairment in AD. The gold standard approach involves clinicopathological investigation, in which the cardinal pathologies are measured post mortem and then correlated with ante mortem measurements of cognitive status. Over 40 clinicopathological studies have been conducted in AD, which have revealed tau-pathology to be the strongest predictor of global cognitive status (Nelson et al., 2012). The relationship between amyloid-pathology and global cognitive measures is relatively weak and attenuated when the effect of tau-pathology is accounted for statistically (Bennett, Schneider, Wilson, Bienias, & Arnold, 2004). The largest, and most recent, clinicopathological study confirmed this pattern, reporting that staging of tau-pathology predicted global cognitive status, while the staging of amyloid-pathology did not (Murray et al., 2015). Neuroimaging evidence has confirmed the contribution of both tau and amyloid to the clinical phenotype, but has been limited in addressing the potential of differential contributions (Bejanin et al., 2017; Hedden, Oh, Younger, & Patel, 2013).

Despite this progress, there remain significant gaps in the literature. First, it is common for markers of clinical status to be reduced to a single score, which is derived from summation across cognitive domains. Relatively few studies have simultaneously investigated the relationship between AD pathology and multiple measures of clinical status. It is possible that the dominance of tau-pathology is not consistently expressed across all clinical symptom domains. Second, a common limitation of clinicopathological research is that histopathological staging is typically recorded using semi-quantitative rating systems (Mirra et al., 1991). While higher scores are associated with greater pathological burden, the intervals between stages are not necessarily equal. Failure to explicitly model the monotonic nature of these variables might have obscured non-linear relationships between pathological staging and measures of clinical severity. Insensitivity to this non-linearity might underpin the negative associations reported for amyloid-pathology. Third, to the best of our knowledge, no studies have directly compared the predictive relationship *between* different pathological staging markers. While the pattern of significant and non-significant findings have been interpreted as differential relationships, true statistical interactions have not been thoroughly investigated (Nieuwenhuis, Forstmann, & Wagenmakers, 2011).

Our study aimed to explicitly address these gaps in the literature by investigating the differential relationship between histopathological staging and multiple clinical severity variables using models that explicitly take into account the ordinal nature of the data. While we expected tau-pathology to be most strongly associated with markers of clinical severity, it was possible that our more appropriate modelling approach would reveal a previously obscured dominant contribution of amyloid.

## Materials and Methods

### Participants

Data for this study were acquired from the National Alzheimer’s Disease Coordinating Center (NACC) neuropathology data set (Besser et al., 2018). This analysis used data from 22 Alzheimer’s Disease Centers (ADCs). Data from visits between September 2005 and November 2017. Participants were included in the study if they: (1) were present in the NACC Neuropathology Data Set, (2) had adequate data in order to classify major neuropathological change, (3) were aged > 50 years of age at time of death, (4) had at least one recorded clinical visit prior to death, (5) had either no vascular pathology or mild vascular pathology on neuropathological examination, (6) had no evidence of Lewy bodies on neuropathological examination, (7) had no evidence of other pathology on neuropathological examination (e.g., FTLD-TDP, FTLD-tau, tangle-only dementia, prion disease, etc.), (8) and spoke English as their primary language.

### Neuropathological staging

Neuropathological staging was performed and recorded as per CERAD protocols (Mirra et al., 1991). Only standard blocks were used to assign histopathological staging as described by Montine and colleagues (2012). The presence of tauopathy was determined using Braak staging for neurofibrillary degeneration (CERAD B score: stage 0 - stage VI; Braak & Braak, 1991). Neuritic plaques were staged according to the density of neocortical plaques with argyrophilic dystrophic neurites (CERAD C score: ‘no neuritic plaques’ to ‘frequent neuritic plaques’). Diffuse plaques were staged according to the density of plaques with non-compact amyloid and no apparent dystrophic features (‘no diffuse plaques’ to frequent diffuse plaques’). We used only the ‘derived’ variables from the NACC database, which ensure consistency over different iterations of the of the neuropathology data set.

### Clinical severity variables

Clinical measurements were obtained from five domains: global cognitive function, dementia stage, functional impairment, neuropsychiatric symptoms, and depressive symptomatology. Global cognitive impairment was measured using the mini mental status examination (MMSE; Folstein, Folstein, & McHugh, 1975), and brief clinician-administered cognitive screening instruments. Possible scores range from 0 to 30 with lower scores indicating greater cognitive impairment. Dementia stage was determined using the CDR® Dementia Staging Instrument (CDR; Morris, 1997) global score, which was used to classify patients as having either no impairment, questionable impairment, mild impairment, moderate impairment, or severe impairment. Functional impairment was determined using the functional activities questionnaire (FAQ; Pfeffer, Kurosaki, Harrah Jr, Chance, & Filos, 1982), a 10-item informant-rated instrument measuring impairments in activities of daily living (e.g., paying bills, preparing a balanced meal, shopping for groceries). Possible scores range from 0 to 30 with greater scores indicating greater impairment. Neuropsychiatric symptoms were measured using the neuropsychiatric inventory questionnaire (NPI-Q; Kaufer et al., 2000), a 12-item informant-report instrument assessing psychiatric symptoms (e.g., delusions, hallucinations, etc…) observed in the last month. The severity of the symptom (1 = *m*i*ld t*o 3 = se*vere)* is recorded along with the resulting carer-distress (0 = *not distressing at all t*o 5 = *extreme or severe distress)*. The total score is the sum of severity and distress scores across all items. Possible scores on the NPI range from 0 to 65, with greater scores indicating greater symptomatology. Depression was assessed using the 15-item geriatric depression scale (GDS; Yesavage et al., 1982), which produces scores ranging from 1 to 15 with greater scores indicating more severe depressive symptomatology.

### Statistical analyses

All statistical analyses were performed in the *R* environment (version 3.6.0; R Core Team, 2019). Bayesian general linear multi-level models (GLMMs) were used to investigate the relationship between pathological staging and clinical variables. The three pathological staging variables were included in each model along with sex, age, and the interval between death and histopathological examination. A random intercept term was included across clinical sites. The following models were specified for continuous clinical severity variables:

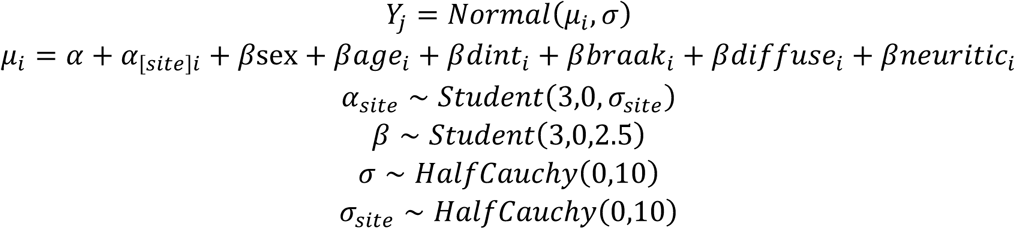

Where *Y*_*j*_ is the continuous severity variable for person *i*. All continuous variables were centred and scaled prior to analysis. The coefficients *βbraak*_*i*_, *βdiffuse*_*i*_, and *βneuritic*_*i*_ were modelled monotonically as:

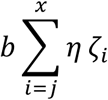

Where *b* determines the size and direction of the effect and *ζ* determines the normalised distances between consecutive predictor categories. A Dirichlet (multivariate beta distribution) prior was placed on the normalised distances with *k* equal to the number of levels of the predictor minus one. Rather than the Gaussian distribution, a cumulative response function was used to model the ordinal structure of the dementia stage (CDR) variable.

All models were estimated using the *brms* package (Bürkner, 2017, 2018). Parameters were estimated via Hamiltonian Markov Chain Monte Carlo (MCMC) implemented in *Stan* (Carpenter et al., 2017) with 3 chains each of 2,000 warm-up and 2,000 actual samples. Successful chain convergence was defined as 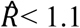 Posterior predictive distributions were inspected to ensure adequate fit and the posterior predictive probability (*PPP*) was computed for each model. Following Gelman and colleagues (2013), .5 < *PPP* < .95 was considered to indicate adequate fit. The Bayesian *R*^2^ was computed to estimate the variance explained by the model in the outcome variable (Gelman, Goodrich, Gabry, & Vehtari, 2018). Following the approach described by McElreath (2018), the difference between predictors was investigated by subtracting samples from the posterior distributions. Point estimates for each parameter are presented as the median of the posterior distribution. Uncertainty around these point estimates is indicated by 95% highest density intervals (HDIs). Sensitivity analyses were performed on subgroups of patients with the time between final clinical visit and death restricted to <=24 months and <= 12 months.

## Results

### Sample characteristics

Of the 144 participants, 71 were female (49%). The mean age at the time of visit was 78.97 (SD = 11.58, min = 47, max = 103). The mean years of education was 15.29 (SD = 3.01, min = 6, max = 20). The mean number of months between final visit and post mortem examination was 14.11 (SD = 13.48, min = 0, max = 66). All participants spoke English as their primary language. The descriptive statistics for clinical outcome variables are shown in *Table 1*.

**Table 1.** Sample characteristics

| *Variable* | *M (SD)* | *Range* |
| --- | --- | --- |
| Age at visit | 78.87 (11.58) | 47 - 103 |
| Global cognition | 19.99 (8.96) | 0 - 30 |
| Functional impairment | 18.86 (12.06) | 0 - 30 |
| Neuropsychiatric symptoms | 9.69 (8.79) | 0 - 38 |
| Depression | 2.83 (2.71) | 0 - 13 |

### Clinical staging

In terms of clinical staging, most participants had a diagnosis of dementia at the time of visit (*n* = 98, 68%). Normal cognition was the next most common diagnosis (*n* = 31, 22%), followed by MCI (*n* = 9, 6%), and ‘impaired but not meeting MCI criteria’ (*n* = 6, 4%). Of those participants classified as having a cognitive impairment, most had a presumed aetiological diagnosis of Alzheimer’s Disease (*n* = 84, 74%). The next most common diagnosis was FTLD (*n* = 13, 12%). The remainder (*n* = 16, 14%) had been diagnosed clinically as either Lewy body disease, coriticobasal degeneration, vascular brain injury, depression, or cognitive impairment due to systemic disease. As noted above, cases confirmed as non-Alzheimer’s disease on postmortem had already been excluded, indicating that the non-Alzheimer clinical diagnoses were not correct.

### Neuropathological staging

The neuropathological staging is summarised in *Table 2*. As shown, while the sample was spread across all pathological stages, the distribution favoured more severe disease.

**Table 2.** Neuropathological staging

| *Pathology* | *Staging* |  |  |  |  |  |  |
| --- | --- | --- | --- | --- | --- | --- | --- |
| *Braak NFT stage* | *0* | *I* | *II* | *III* | *IV* | *V* | *VI* |
| *n (%)* | 4 (3) | 13 (9) | 16 (11) | 19 (13) | 26 (18) | 23 (16) | 43 (30) |
| *Diffuse plaques* | *No plaques* | *Sparse* | *Moderate* | *Frequent* |  |  |  |
| *n (%)* | 19 (13) | 29 (20) | 23 (16) | 73 (51) |  |  |  |
| *Neuritic plaques* | *No plaques* | *Sparse* | *Moderate* | *Frequent* |  |  |  |
| *n (%)* | 28 (19) | 15 (10) | 31 (22) | 70 (49) |  |  |  |

### Predictors of clinical status

Separate Bayesian GLMMs were computed for each clinical severity variable. Examination of the posterior predictive distributions and residuals revealed adequate model fit (PPPs = .32 −.59). The posterior predictive distributions are included in *Figure S1*. As shown in *Table 3*, the models explained a moderate amount of variance in global cognition, functional impairment, and neuropsychiatric symptoms (*R*^2^ = .46 − .58). The model was comparatively poor at explaining variance in depression (*R*^2^ = .14).

**Table 3.** Model coefficients for neuropathological staging and clinical outcome

| *Outcome* | *n* | *PPP* | *R*<sup>2</sup> | *Braak NFT stage* | *Diffuse plaques* | *Neuritic plaques* |
| --- | --- | --- | --- | --- | --- | --- |
| Global cognition | 111 | .59 | .56 | **-1.46**<br>[-2.14, -0.86] | 0.22<br>[-0.36, 0.75] | -0.43<br>[-0.99, 0.07] |
| Dementia stage | 144 | .41 | - | **5.45**<br>[3.73, 7.50] | 0.27<br>[-0.52, 0.94] | 0.05<br>[-0.59, 0.70] |
| Functional impairment | 116 | .51 | .58 | **1.46**<br>[0.90, 2.06] | 0.27<br>[-0.52, 0.94] | 0.05<br>[-0.59, 0.70] |
| Neuropsychiatric symptoms | 136 | .43 | .46 | **1.13**<br>[0.54, 1.78] | -0.05<br>[-0.54, 0.48] | -0.18<br>[-0.74, 0.40] |
| Depression | 100 | .32 | .14 | -0.11<br>[-1.10, 0.86] | 0.33<br>[-0.38, 1.09] | **-0.89**<br>[-1.62, -0.14] |

**Note:** *n* = number of participants included in analysis following list-wise removal of cases with missing outcome data. *PPP* = posterior predictive probability. *R*<sup>2</sup> = Bayesian coefficient of determination (not computed for dementia stage due to cumulative response function). All parameters are the median of the posterior distribution [95% highest density intervals].

As shown in *Figure 1*, Braak NFT staging was a significant predictor of all clinical severity variables except for depression. Higher Braak NFT stage was associated with worse cognitive function (β = −1.46 [−2.14, −0.86]), later dementia stage (β = 5.45 [3.73, 7.50]), greater functional impairment (β = 1.46 [0.90, 2.06]), and more severe neuropsychiatric symptoms (β = 1.13 [0.54, 1.78]). There was no evidence for a relationship between diffuse plaque stage and any clinical severity variable. Higher neuritic plaque stage was associated with lower depressive symptomatology (β = −0.89 [−1.62, −0.14]). No other relationships between clinical variables and neuritic plaque stage were supported. The relationship between Braak NFT stage and clinical severity is shown in *Figure 2* and the associations with all neuropathological variables is show in *Figure S2*. Full model coefficients (including nuisance parameters) are included in *Table S2*.

**Figure 1.**
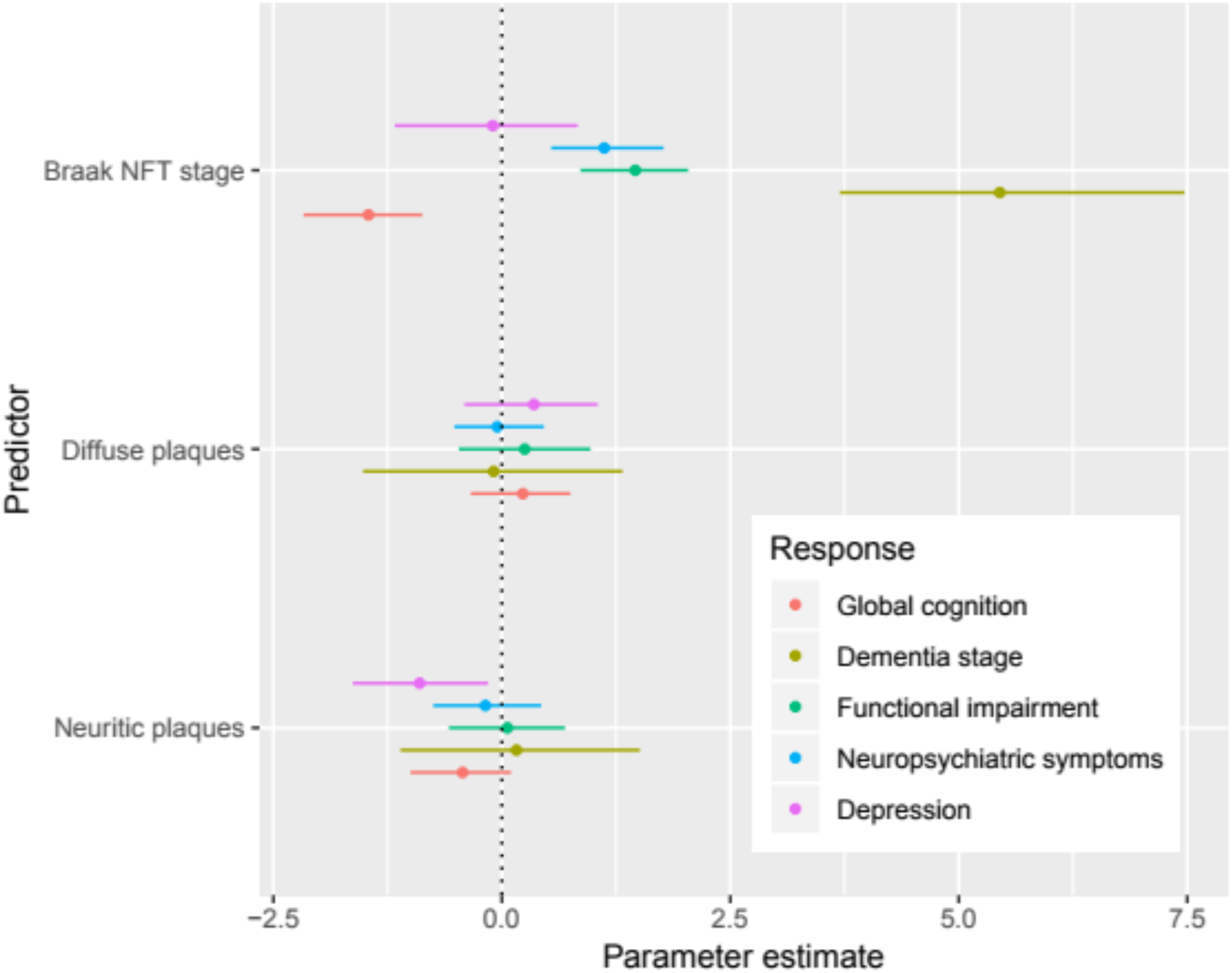
The coefficients for all five models of clinical severity outcomes. Parameters are shown as the median of the posterior distribution with 95% highest posterior density intervals (HDPIs). As show, greater Braak NFT stage was associated with worse cognition, later dementia stage, greater functional impairment, and more neuropsychiatric symptoms. No other predictors were statistically supported, except for a slight effect of neuritic plaque stage on depression.

**Figure 2.**
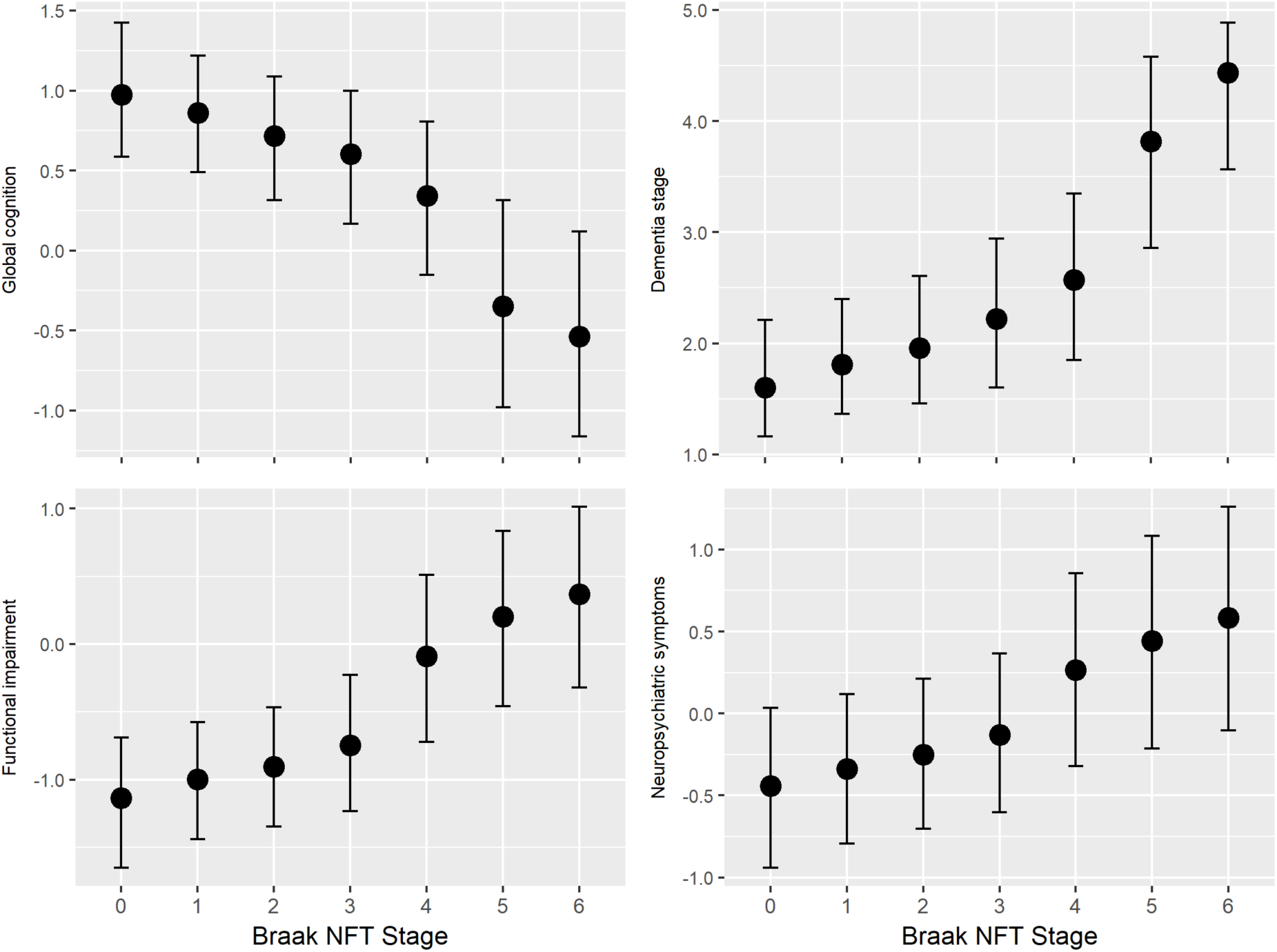
The marginal effects plots for NFT stage show strong relationships between pathological staging and clinical variables. Higher Braak staging was associated with more impaired global cognition, greater functional impairment, more neuropsychiatric symptoms, and later dementia stage. The relationships are slightly non-linear, showing acceleration at later Braak stages.

### Difference between predictors

The relative predictive contribution of Braak NFT stage compared to diffuse and neuritic plaques was examined by computing the posterior probability distribution of the difference between the coefficients (as described in methods section above). The results from this analysis are shown in *Table 4*. When directly compared to diffuse or neuritic plaques, Braak NFT stage was a stronger predictor of global cognition, dementia stage, functional impairment, and neuropsychiatric symptoms. The evidence did not support such differences for depression.

**Table 4.** Comparison between coefficients – Median [95% HDIs]

| *Outcome* | *Braak NFT Stage*<br>*> diffuse plaques* | *Braak NFT Stage*<br>*> neuritic plaques* |
| --- | --- | --- |
| Global cognition | **1.28 [0.63, 1.92]** | **1.16 [0.20, 1.99]** |
| Dementia stage | **5.37 [3.27, 7.71]** | **5.44 [3.20, 7.79]** |
| Functional impairment | **1.14 [0.26, 1.93]** | **1.26 [0.51, 1.93]** |
| Neuropsychiatric symptoms | **0.85 [0.16, 1.47]** | **0.79 [0.13, 1.38]** |
| Depression | -0.01 [-0.72, 0.76] | -0.31 [-1.13, 0.77] |

### Sensitivity analysis

The time between clinical assessment and post-mortem varied significantly. To examine robustness of the above findings, all analyses were repeated on subgroup of patients who underwent post-mortem examination within 24 months of the final clinical visit (*n* = 118, 82% of the sample). A second sensitivity analysis was then performed on patients who died within 12 months of their final visit (*n* = 84, 58% of the sample). These results are shown in *Table S2*. The relationships between Braak NFT stage and clinical variables were replicated in the 24-month subgroup. As shown in *Table S3*, the relationships between Braak NFT stage and global cognition, dementia stage, and functional impairment were also replicated in the 12-month subgroup, while the relationship with neuropsychiatric symptoms was not. The relationship between neuritic plaques and depressive symptomatology was not replicated in either subgroup analysis. Otherwise the findings for neuritic and diffuse plaques were comparable to the primary analysis.

## Discussion

In this study, we investigated the relationship between clinical symptom severity and AD neuropathological staging confirmed via post-mortem examination. Braak NFT staging was associated with global cognitive impairment, dementia stage, functional impairment, and neuropsychiatric symptoms. In contrast, the pathological staging of diffuse and neuritic plaques was not associated any of these clinical variables. Direct comparison between the different pathological markers confirmed that Braak NFT staging had a stronger relationship to the clinical severity variables compared to neuritic and diffuse plaques. The advanced statistical modelling techniques used for this study mean that the findings are robust, in part because we explicitly modelled the ordinal nature of neuropathological staging data. To the best of our knowledge, our study was the first to model the clincopathological nexus in Alzheimer’s disease using such a statistical approach.

Overall, our findings are consistent with previous clinicopathological investigations in Alzheimer’s disease. For example, in a comprehensive review of the literature, Nelson and colleagues (Nelson et al., 2012) also concluded that tau-related pathology had a stronger relationship with clinical variables compared to amyloid-related pathology. Similar findings were reported by more recent studies (Bennett et al., 2004; Murray et al., 2015). Importantly, our findings do not suggest that amyloid-pathology is not related to clinical severity. Rather, these findings suggest that when amyloid and tau-pathology are considered together, tau-pathology is the driving factor behind clinical impairment. This is an important consideration, as tau and amyloid-pathology are highly collinear which means that examination the relationship between clinical severity and either pathology in isolation is likely to always reveal positive correlations. It is only by examining the two pathologies are included in a single model that the differential relationship emerges.

Our findings are also consistent with previous work investigating the relationship between clinical severity and non-histopathological biomarkers. For example, a great number of studies have shown correlations between amyloid-burden revealed by imaging with PET ligands (Bejanin et al., 2017; Hedden et al., 2013). The in vivo imaging of tau-related pathology is relatively less developed, but has nonetheless revealed correlations between pathological burden and clinical staging (Jack et al., 2018; Johnson et al., 2016; Pontecorvo et al., 2017).

Our findings suggest that the dominance of tau-related pathology extends across different aspects of the clinical phenotype, including dementia staging, global cognitive impairment, functional disability, and neuropsychiatric symptoms. Depressive symptomatology, however, was not convincingly associated with one pathology or another. This builds on previous work by our group demonstrating a differential relationship between pathology, cognitive, and network disruption. For example, we have shown that memory is most strongly associated with cerebrospinal fluid markers of tau-pathology while amyloid-pathology is preferentially associated with diffusely represented neurocognitive functions such as processing speed (Malpas et al., 2015). We have also demonstrated a preferential relationship between tau-pathology and mesial temporal functional connectivity (Malpas, Saling, Velakoulis, Desmond, & O’Brien, 2016). Amyloid, on the other hand, appears to preferentially affect structures outside the mesial temporal regions (Malpas et al., 2018).

The present study had a number of strengths. First, for a clinicopathological study, our sample consistent of a large number of patients. Second, the advanced analytical methods represent a significant improvement over previous studies in this area. By using multi-variable models, we were able to examine the relationship between clinical markers and different neuropathological staging variables simultaneously. In other words, we were able to examine the effect of one staging variable (e.g., Braak stage) while holding the other staging variables constant. Unlike dominant univariate analyses, which only consider one pathological marker at a time, our approach allowed for the relative contribution of each pathological marker to be investigated. Third, our study was also novel in that we explicitly modelled the ordinal nature of neuropathological staging data. The typical approach to analyses this sort of data is to treat the variables as continuous (which violates model assumptions), or to consider it to be nominal (i.e., categorical) which reduces statistical sensitivity. By modelling these variables as monotonic functions we were able to investigate the overall direction of the effect via while acknowledging the ordinal nature of the underlying relationship. comparing the difference Fourth, the key results survived sensitivity analyses that addressed the issue of variable time between last clinical visit and post-mortem analysis. Fifth, by directly between model parameters we were able to convincingly show that Braak NFT staging was a stronger predictor compared to other pathological staging variables. Rather than just showing that one predictor was statistically significant, while the others were not, our approach allowed us to demonstrate an outcome by predictor interaction (Nieuwenhuis et al., 2011). Finally, our use of multiple outcome variables provides evidence that the different in underlying pathological staging has implications not only for cognition, but a range of other phenotype markers, including functional impairment and neuropsychiatric symptoms.

Despite these strengths, some limitations must be considered. First, while we took a multi-construct approach to examining clinical severity, the instrument used to assess global cognitive function suffers from the problem of summation. Specifically, the MMSE adds a number of cognitive tasks together for form a single score (Folstein et al., 1975). As a summative measure, it is therefore not possible to investigate whether specific neuropathological staging variable had preferential effects on different aspects of the cognitive phenotype. As noted above, previous work by our group has suggested that tau-and amyloid-related pathology might have differential effects on cognition that could not be confirmed in this study (Malpas et al., 2015).

Given the advent of reliable in vivo imaging of both tau- and amyloid-related pathology, a clear direction of future work is to fully examine the preferential clinic-pathological contributions of tau and amyloid (Villemagne, Fodero-Tavoletti, Masters, & Rowe, 2015). This should not only examine the difference is magnitude of the relationship, but also consider whether specific cognitive domains are indeed affected preferentially by each of the pathologies. Given that therapeutic trials tend to target either tau or amyloid, but not both, it will be essential to choose neurocognitive outcome measures that accurately target the relevant pathological system. Failure to take into account the preferential contribution of

In conclusion, we investigated the relationship between three histopathological staging variables and markers of clinical severity in patients with and without cognitive impairment. These findings suggest that tau-pathology, in the form of Braak NFT staging, is a strong predictor of global cognitive status, clinical dementia stage, functional abilities, and neuropsychiatric symptoms. Amyloid-pathology, in the form of neuritic and diffuse plaques, was not independently associated with any markers of clinical impairment. These findings confirm previous clinicopathological studies and suggest that the previously reported dominance of tau-pathology is not merely an artefact of the ordinal semi-quantitative staging systems. These findings underscore the need to match clinical markers to the relevant pathological substrate in clinical trials. Outcome measures that heavily weight cognitive, functional, or neuropsychiatric symptoms are likely to be insensitive to interventions that target amyloid-pathology in isolation.

## Acknowledgements

The authors declare no conflicts of interest.

The NACC database is funded by NIA/NIH Grant U01 AG016976. NACC data are contributed by the NIA-funded ADCs: P30 AG019610 (PI Eric Reiman, MD), P30 AG013846 (PI Neil Kowall, MD), P50 AG008702 (PI Scott Small, MD), P50 AG025688 (PI Allan Levey, MD, PhD), P50 AG047266 (PI Todd Golde, MD, PhD), P30 AG010133 (PI Andrew Saykin, PsyD), P50 AG005146 (PI Marilyn Albert, PhD), P50 AG005134 (PI Bradley Hyman, MD, PhD), P50 AG016574 (PI Ronald Petersen, MD, PhD), P50 AG005138 (PI Mary Sano, PhD), P30 AG008051 (PI Thomas Wisniewski, MD), P30 AG013854 (PI M. Marsel Mesulam, MD), P30 AG008017 (PI Jeffrey Kaye, MD), P30 AG010161 (PI David Bennett, MD), P50 AG047366 (PI Victor Henderson, MD, MS), P30 AG010129 (PI Charles DeCarli, MD), P50 AG016573 (PI Frank LaFerla, PhD), P50 AG005131 (PI James Brewer, MD, PhD), P50 AG023501 (PI Bruce Miller, MD), P30 AG035982 (PI Russell Swerdlow, MD), P30 AG028383 (PI Linda Van Eldik, PhD), P30 AG053760 (PI Henry Paulson, MD, PhD), P30 AG010124 (PI John Trojanowski, MD, PhD), P50 AG005133 (PI Oscar Lopez, MD), P50 AG005142 (PI Helena Chui, MD), P30 AG012300 (PI Roger Rosenberg, MD), P30 AG049638 (PI Suzanne Craft, PhD), P50 AG005136 (PI Thomas Grabowski, MD), P50 AG033514 (PI Sanjay Asthana, MD, FRCP), P50 AG005681 (PI John Morris, MD), P50 AG047270 (PI Stephen Strittmatter, MD, PhD).

## References

Arnold, S. E., Hyman, B. T., Flory, J., Damasio, A. R., & Van Hoesen, G. W. (1991). The topographical and neuroanatomical distribution of neurofibrillary tangles and neuritic plaques in the cerebral cortex of patients with Alzheimer’s disease. Cerebral cortex, 1(1), 103–116.

Arriagada, P. V., Growdon, J. H., Hedley-Whyte, E. T., & Hyman, B. T. (1992). Neurofibrillary tangles but not senile plaques parallel duration and severity of Alzheimer’s disease. Neurology, 42(3), 631–631.

Bejanin, A., Schonhaut, D. R., La Joie, R., Kramer, J. H., Baker, S. L., Sosa, N., … Lauriola, M. (2017). Tau pathology and neurodegeneration contribute to cognitive impairment in Alzheimer’s disease. Brain, 140(12), 3286–3300.

Bennett, D. A., Schneider, J. A., Wilson, R. S., Bienias, J. L., & Arnold, S. E. (2004). Neurofibrillary tangles mediate the association of amyloid load with clinical Alzheimer disease and level of cognitive function. Archives of neurology, 61(3), 378–384.

Besser, L. M., Kukull, W. A., Teylan, M. A., Bigio, E. H., Cairns, N. J., Kofler, J. K., … Nelson, P. T. (2018). The revised National Alzheimer’s Coordinating Center’s Neuropathology Form—available data and new analyses. Journal of Neuropathology & Experimental Neurology, 77(8), 717–726.

Braak, H., Alafuzoff, I., Arzberger, T., Kretzschmar, H., & Del Tredici, K. (2006). Staging of Alzheimer disease-associated neurofibrillary pathology using paraffin sections and immunocytochemistry. Acta neuropathologica, 112(4), 389–404.

Braak, H., & Braak, E. (1991). Neuropathological stageing of Alzheimer-related changes. Acta neuropathologica, 82(4), 239–259.

Bürkner, P.-C. (2017). brms: An R Package for Bayesian Multilevel Models Using Stan. Journal of Statistical Software, 80(1), 1–28. doi:10.18637/jss.v080.i01

Bürkner, P.-C. (2018). Advanced Bayesian Multilevel Modeling with the R Package brms. The R Journal, 10(1), 395–411. doi:10.32614/RJ-2018-017

Carpenter, B., Gelman, A., Hoffman, M. D., Lee, D., Goodrich, B., Betancourt, M., … Riddell, A. (2017). Stan: A probabilistic programming language. Journal of Statistical Software, 76(1).

Cho, H., Choi, J. Y., Hwang, M. S., Kim, Y. J., Lee, H. M., Lee, H. S., … Lyoo, C. H. (2016). In vivo cortical spreading pattern of tau and amyloid in the Alzheimer disease spectrum. Annals of neurology, 80(2), 247–258.

Folstein, M. F., Folstein, S. E., & McHugh, P. R. (1975). “Mini-mental state”: a practical method for grading the cognitive state of patients for the clinician. Journal of psychiatric research, 12(3), 189–198.

Gelman, A., Goodrich, B., Gabry, J., & Vehtari, A. (2018). R-squared for Bayesian regression models. The American Statistician(just-accepted), 1–6.

Gelman, A., Stern, H. S., Carlin, J. B., Dunson, D. B., Vehtari, A., & Rubin, D. B. (2013). Bayesian data analysis: Chapman and Hall/CRC.

Haass, C., & Selkoe, D. J. (2007). Soluble protein oligomers in neurodegeneration: lessons from the Alzheimer’s amyloid β-peptide. Nature reviews Molecular cell biology, 8(2), 101.

Hedden, T., Oh, H., Younger, A. P., & Patel, T. A. (2013). Meta-analysis of amyloid-cognition relations in cognitively normal older adults. Neurology, 80(14), 1341–1348.

Jack, C. R., Wiste, H. J., Schwarz, C. G., Lowe, V. J., Senjem, M. L., Vemuri, P., … Gunter, L. (2018). Longitudinal tau PET in ageing and Alzheimer’s disease. Brain, 141(5), 1517–1528.

Johnson, K. A., Schultz, A., Betensky, R. A., Becker, J. A., Sepulcre, J., Rentz, D., … Papp, (2016). Tau positron emission tomographic imaging in aging and early A lzheimer disease. Annals of neurology, 79(1), 110–119.

Kaufer, D. I., Cummings, J. L., Ketchel, P., Smith, V., MacMillan, A., Shelley, T., … DeKosky, S. T. (2000). Validation of the NPI-Q, a brief clinical form of the Neuropsychiatric Inventory. The Journal of neuropsychiatry and clinical neurosciences, 12(2), 233–239.

Malpas, C. B., Saling, M. M., Velakoulis, D., Desmond, P., Hicks, R. J., Zetterberg, H., … O’Brien, T. J. (2018). Cerebrospinal Fluid Biomarkers are Differentially Related to Structural and Functional Changes in Dementia of the Alzheimer’s Type. Journal of Alzheimer’s Disease, 62(1), 417–427.

Malpas, C. B., Saling, M. M., Velakoulis, D., Desmond, P., & O’Brien, T. J. (2016). Differential functional connectivity correlates of cerebrospinal fluid biomarkers in dementia of the Alzheimer’s type. Neurodegenerative Diseases, 16(3-4), 147–151.

Malpas, C. B., Saling, M. M., Velakoulis, D., Desmond, P., & O’Brien, T. J. (2015). Tau and amyloid-β cerebrospinal fluid biomarkers have differential relationships with cognition in mild cognitive impairment. Journal of Alzheimer’s Disease, 47(4), 965–975.

McElreath, R. (2018). Statistical rethinking: A Bayesian course with examples in R and Stan: Chapman and Hall/CRC.

Mirra, S. S., Heyman, A., McKeel, D., Sumi, S., Crain, B. J., Brownlee, L., … Berg, L. (1991). The Consortium to Establish a Registry for Alzheimer’s Disease (CERAD): Part II. Standardization of the neuropathologic assessment of Alzheimer’s disease. Neurology, 41(4), 479–479.

Montine, T. J., Phelps, C. H., Beach, T. G., Bigio, E. H., Cairns, N. J., Dickson, D. W., … Mirra, S. S. (2012). National Institute on Aging–Alzheimer’s Association guidelines for the neuropathologic assessment of Alzheimer’s disease: a practical approach. Acta neuropathologica, 123(1), 1–11.

Morris, J. C. (1997). Clinical dementia rating: a reliable and valid diagnostic and staging measure for dementia of the Alzheimer type. International psychogeriatrics, 9(S1), 173–176.

Murray, M. E., Lowe, V. J., Graff-Radford, N. R., Liesinger, A. M., Cannon, A., Przybelski, S. A., … Kantarci, K. (2015). Clinicopathologic and 11C-Pittsburgh compound B implications of Thal amyloid phase across the Alzheimer’s disease spectrum. Brain, 138(5), 1370–1381.

Nelson, P. T., Alafuzoff, I., Bigio, E. H., Bouras, C., Braak, H., Cairns, N. J., … Tredici, K. D. (2012). Correlation of Alzheimer disease neuropathologic changes with cognitive status: a review of the literature. Journal of Neuropathology & Experimental Neurology, 71(5), 362–381.

Nieuwenhuis, S., Forstmann, B. U., & Wagenmakers, E.-J. (2011). Erroneous analyses of interactions in neuroscience: a problem of significance. Nature neuroscience, 14(9), 1105.

Pfeffer, R., Kurosaki, T., Harrah Jr, C., Chance, J., & Filos, S. (1982). Measurement of functional activities in older adults in the community. Journal of gerontology, 37(3), 323–329.

Pontecorvo, M. J., Devous Sr, M. D., Navitsky, M., Lu, M., Salloway, S., Schaerf, F. W., … Lim, N. C. (2017). Relationships between flortaucipir PET tau binding and amyloid burden, clinical diagnosis, age and cognition. Brain, 140(3), 748–763.

R Core Team. (2019). R: A Language and Environment for Statistical Computing. Vienna, Austria: R Foundation for Statistical Computing.

Thal, D. R., Rüb, U., Orantes, M., & Braak, H. (2002). Phases of Aβ-deposition in the human brain and its relevance for the development of AD. Neurology, 58(12), 1791–1800.

Villemagne, V. L., Fodero-Tavoletti, M. T., Masters, C. L., & Rowe, C. C. (2015). Tau imaging: early progress and future directions. The Lancet Neurology, 14(1), 114–124.

Yesavage, J. A., Brink, T. L., Rose, T. L., Lum, O., Huang, V., Adey, M., & Leirer, V. O. (1982). Development and validation of a geriatric depression screening scale: a preliminary report. Journal of psychiatric research, 17(1), 37–49.

