## Supplemental Information for "The histopathological staging of tau, but not amyloid, corresponds to antemortem cognitive status, dementia stage, functional abilities, and neuropsychiatric symptoms"

### Supplemental Materials

*Table S1.* All model coefficients for the full sample. Coefficients are medians of the posterior distribution [95% highest density intervals].

| *Parameter* | *Global cognition* | *Dementia stage* | *Functional impairment* | *Neuropsychiatric symptoms* | *Depression* |
|---|---|---|---|---|---|
| *Coefficients* | | | | | |
| Sex | -0.09 [-0.38, 0.19] | 0.12 [-0.58, 0.81] | 0.09 [-0.18, 0.37] | -0.13 [-0.41, 0.14] | 0.27 [-0.13, 0.69] |
| Age | **0.25 [0.06, 0.41]** | -0.30 [-0.73, 0.14] | **-0.20 [-0.36, -0.06]** | **-0.27 [-0.43, -0.10]** | -0.09 [-0.30, 0.14] |
| Death interval | **0.19 [0.04, 0.35]** | **-0.69 [-1.08, -0.27]** | **-0.16 [-0.30, -0.01]** | **-0.21 [-0.35, -0.05]** | -0.10 [-0.33, 0.12] |
| Braak NFT stage | **-1.46 [-2.14, -0.86]** | **5.45 [3.73, 7.50]** | **1.46 [0.90, 2.06]** | **1.13 [0.54, 1.78]** | -0.11 [-1.10, 0.86] |
| Diffuse plaques | 0.22 [-0.36, 0.75] | -0.07 [-1.47, 1.36] | 0.27 [-0.52 , 0.94] | -0.05 [-0.54, 0.48] | 0.33 [-0.38, 1.09] |
| Neuritic plaques | -0.43 [-0.99, 0.07] | 0.16 [-1.12, 1.51] | 0.05 [-0.59 , 0.70] | -0.18 [-0.74, 0.40] | **-0.89 [-1.62, -0.14]** |
| *Model fit* | | | | | |
| *n* | 111 | 144 | 116 | 136 | 100 |
| *PPP* | .50 | .42 | .51 | .50 | .49 |
| $R^2$ | .56 | - | .58 | .46 | .18 |

**Note:** *n* = number of participants included in analysis following list-wise removal of cases with missing outcome data. *PPP* = posterior predictive probability. $R^2$ = Bayesian coefficient of determination (not computed for dementia stage due to cumulative response function). All parameters are the median of the posterior distribution [95% highest density intervals].

*Table S2.* Model coefficients limited to patients with cognitive examination <= 24 months prior to death.

| *Outcome* | *n* | *PPP* | $R^2$ | *Braak NFT stage* | *Diffuse plaques* | *Neuritic plaques* |
|---|---|---|---|---|---|---|
| Global cognition | 92 | .63 | .60 | **-1.45 [-1.96, -0.97]** | 0.13 [-0.36, 0.61] | -0.47 [-0.95, 0.02] |
| Dementia stage | 118 | .45 | - | **5.09 [3.24, 7.21]** | -0.04 [-1.58, 1.43] | 0.58 [-0.91, 2.03] |
| Functional impairment | 95 | .49 | .56 | **1.31 [0.68, 1.94]** | -0.05 [-0.73, 0.77] | 0.35 [-0.44, 1.02] |
| Neuropsychiatric symptoms | 110 | .44 | .49 | **1.00 [0.30, 1.72]** | -0.06 [-0.58, 0.48] | -0.04 [-0.70, 0.63] |
| Depression | 81 | .36 | .22 | 0.00 [-1.04, 1.00] | 0.14 [-0.64, 0.89] | -0.65 [-1.40, 0.32] |

**Note:** *n* = number of participants included in analysis following list-wise removal of cases with missing outcome data. *PPP* = posterior predictive probability. $R^2$ = Bayesian coefficient of determination (not computed for dementia stage due to cumulative response function). All parameters are the median of the posterior distribution [95% highest density intervals].

*Table S3.* Model coefficients limited to patients with cognitive examination <= 12 months prior to death.

| *Outcome* | *n* | *PPP* | $R^2$ | *Braak NFT stage* | *Diffuse plaques* | *Neuritic plaques* |
|---|---|---|---|---|---|---|
| Global cognition | 64 | .49 | .59 | **-1.29 [-1.94, -0.63]** | 0.11 [-0.42, 0.65] | -0.54 [-1.16 , 0.01] |
| Dementia stage | 84 | .52 | - | **4.84 [2.37, 7.32]** | 0.25 [-1.41, 1.97] | 0.50 [-1.30, 2.35] |
| Functional impairment | 70 | .51 | .61 | **1.35 [0.66, 2.06]** | -0.23 [-0.82, 0.45] | 0.22 [-0.56, 0.94] |
| Neuropsychiatric symptoms | 80 | .50 | .45 | 0.63 [-0.35, 1.55] | -0.12 [-0.80, 0.53] | 0.17 [-0.68, 0.99] |
| Depression | 58 | .50 | .26 | -0.27 [-1.49, 0.98] | 0.17 [-0.71, 1.06] | -0.67 [-1.60, 0.42] |

**Note:** *n* = number of participants included in analysis following list-wise removal of cases with missing outcome data. *PPP* = posterior predictive probability. $R^2$ = Bayesian coefficient of determination (not computed for dementia stage due to cumulative response function). All parameters are the median of the posterior distribution [95% highest density intervals].

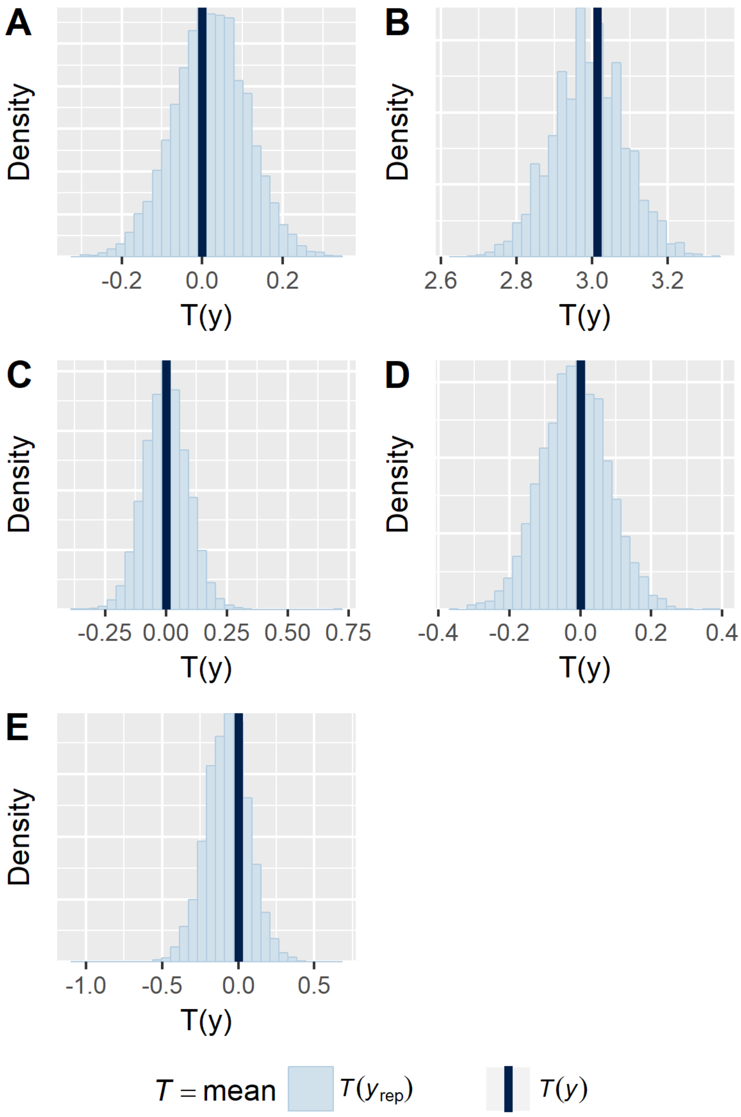


*Figure S1.* Posterior probability distribution checks. **A** = global cognition, **B** = dementia stage, **C** = functional impairment, **D** = neuropsychiatric symptoms, **E** = depression. The solid blue bar, T(y), shows the mean of the observed response variable. Blue bars represent the distribution of means of response variable for each sample in the posterior predictive distribution. All models produced symmetric distributions around the observed mean. Posterior predictive probability values are presented in the body of the manuscript.

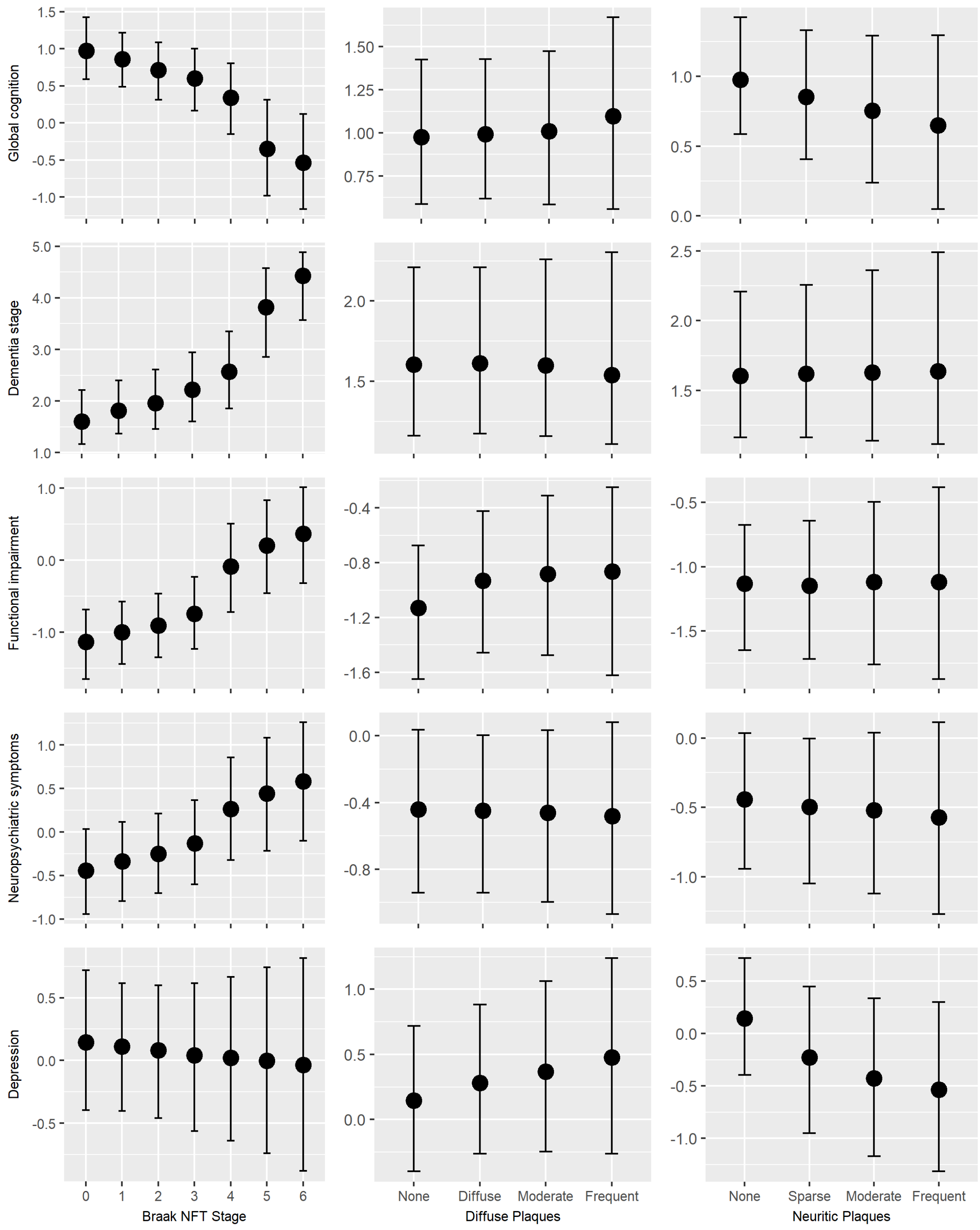


*Figure S2.* Marginal effects across pathological staging predictors.
